# Hygienic behaviour selection via freeze-killed honey bee brood not associated with chalkbrood resistance in eastern Australia

**DOI:** 10.1101/407338

**Authors:** Jody Gerdts, R. Laurie Dewar, Michael Simone-Finstrom, Trevor Edwards, Michael Angove

**Author notes:** Corresponding Author (JG). Current Address: Department of Pharmacy and Applied Science La Trobe University, Edwards Road, Flora Hill Victoria, 3552. These authors contributed equally to this work (R. Laurie Dewar, Michael Simone-Finstrom). These authors also contributed equally to this work (Trevor Edwards, Michael Angove).

## Abstract

Hygienic behaviour is a social immune response in honey bees shown to help provide resistance to honey bee pests and diseases. A survey of hygienic behaviour and brood diseases was conducted on 649 colonies in eastern Australia to initiate a selective breeding program targeting disease resistance and provide a level of resistance to Varroa (*Varroa destructor* Anderson and Trueman and *V. jacobsoni* Oudemans) mites should they become established in Australia. The test population showed a remarkably high baseline level of hygienic behaviour with 17% of colonies meeting or exceeding breeding selection thresholds. Colonies belonging to a breeding program were 5.8 times more likely to be highly hygienic and colonies headed by queens raised from hygienic queen mothers were 2.2 times more likely. Nectar availability (nectar yielding flowering plants within honey bee forage range) influenced hygienic behaviour expression but was not a significant predictor of level of hygienic behaviour. Surprisingly, hygienic behaviour was not a significant predictor of the presence of infection of the honey bee brood disease chalkbrood (*Ascosphaera apis*) and was not influential in predicting severity of chalkbrood the infection in surveyed honey bee colonies. This study, along with reports from commercial beekeepers that chalkbrood infection is on the rise, warrants a deeper exploration of the host-pathogen relationship between *Apis mellifera* and *Ascosphaera apis* in Australia.

## Introduction

European honey bees (Apis mellifera), are an integral component of food security worldwide. In Australia, honey bees provide essential pollination services for food production, contributing directly to between $8.35 billion and $19.97 billion worth of horticultural and agricultural production annually and produce between 20,000 and 30,000 tons of honey a year valued at $90 million [1]. Honey bee contribution to Australian food security is only possible because of limited pest and disease exposure as a direct consequence of the country’s isolation. Although several major brood diseases are endemic such as, American Foulbrood (*Paenibacillus larvae*) European Foulbrood (*Melissococcus pluton*), and chalkbrood (Ascosphaera apis), Australia remains the only country with a significant apicultural industry free of *Varroa* mites (*Varroa destructor* Anderson and Trueman and *V. jacobsoni* Oudemans).

*V. destructor* is the foremost cause of honey bee colony losses globally [2,3], impacting both pollination services and honey production. *Varroa* parasitism has been shown to vector deadly viruses [4] while chemical treatments used to control *Varroa* in honey bee hives can affect reproduction [5,6,7] and interact negatively with other agricultural chemicals [8]. Furthermore, *Varroa* species have been shown to develop resistance to acaricides [9,10] rendering the reliance on these chemicals unsustainable in the long term.

Once established, the mite will have devastating consequences for Australia’s honey bee reliant industries and will progressively destroy most to all feral colonies [11,12,13]. This will place an unprecedented demand on apiarists for managed colonies for pollination. In the 1980s *Varroa* decimated honey bees in the United States and in the early 2000s caused similar collapse in bee populations in New Zealand [14]. Currently, much of Australian’s pollination services are performed by feral honey bee colonies. Reliance on this incidental pollination understates the role that professional apiculturists will need to play in national food security particularly in light of this potential parasite threat.

One strategy to enhance the resilience of Australian honey bee stocks is to breed for behavioural traits of resistance to various pathogens and parasites; hygienic behaviour is one such trait. Hygienic behaviour is a form of social immunity [15] and has been studied extensively over the past 80 years [16,17]. This heritable behaviour imparts a specific type of pathogen defense triggered by olfactory cues [18] where 15-18 day-old workers [19] detect, uncap, and remove dead or diseased brood from the nest before the disease enters an infectious phase or the pest (*Varroa*) has a chance to reproduce. Consequently, bees bred for hygienic behavior are reported to have a natural resistance to microbial diseases [20] such as chalkbrood [21], American foulbrood [22] and viruses [23], and also to non-microbial agents, such as *Varroa* mites [24,25]. Selecting for hygienic behaviour in colonies through specific breeding processes has yet to show any negative colony level effects [26] or individual [27] trade-offs, and has been employed by breeding programs across the world to enhance colony social immunity.

Research over the past two decades has shown that hygienic behavioural traits are present in Australian honey bees [28, 29] providing the opportunity to select from honey bee populations adapted to Australia’s unique environment and flora. Meixner and colleagues [30] demonstrated that disease resistance and *Varroa* tolerance are closely tied to adaptations linked with specific environmental cues and when those cues are absent, disease resistance and pest tolerance significantly decrease. It is therefore pre-emptive to selectively breed these traits into stock suited to Australian conditions; building resilience and providing resistance to endemic diseases, and preparing for living with *Varroa* and associated viruses.

In order to commence selective breeding for hygienic behaviour, we undertook a large-scale assessment of 649 colonies across 9 beekeeping operations to generate a working baseline of the level of hygienic behaviour in Australian production and breeding colonies. During this assessment, we recorded the prevalence of brood diseases affecting the colonies and if the bees has access to nectar yielding flower (referred to as ‘nectar flow’). Our goals were to identify potential hygienic breeding stock, determine if queen selection alone was sufficient to confer hygienic behaviour, assess the influence of nectar flow on the expression of hygienic behaviour, and gauge the efficacy of hygienic behaviour using chalkbrood presence as an indicator.

## Materials and Methods

From late April (early autumn) 2014 to February 2016 (late summer), commercial production colonies and colonies containing breeding lines (N=649) from 9 beekeeping operations in 21 bee yards from eastern Australia (Tasmania, Victoria, New South Wales, and Queensland) were assessed for hygienic behavior. Three operations conducted selective breeding programs while six operations were primarily geared toward pollination and honey production. The freeze-kill brood (FKB) assay [31] was used on 50-100 colonies per operation and ranged in size from nucleus colonies (4 frames of bees and 2-3 frames of brood) to honey production colonies (16+ frames of bees and 6-8 frames of brood). Colonies tested met key selection traits (eg. honey production, temperament, etc) or belonged to specific breeding lines. Whether or not the queen heading the colony was a daughter of a hygienic mother (referred to as ‘queen selection’) and if the colony belonged to a breeding program was noted. The presence or absence of chalkbrood disease and the presence of a nectar flow was also recorded. When a colony was infected with chalkbrood disease, the severity of the disease was rated on a 1-3 scale as determined by the number of mummies or infected larvae on both sides of the middle brood frame: slight 1-10, moderate 11-40, severe 40+.

Although hygienic behaviour can be scored as a continuum from 0-100, represented by a percentage of dead brood removed by nurse bees in 24 hours, it is common practice to classify colonies on a binary scale as hygienic or not hygienic using 95% removal rates as the categorization threshold [32,33]. Strict hygienic behaviour (score based only on full removal of killed brood) is frequently used for selective breeding purposes while liberal hygienic behaviour (score based on a combination of full and partial removal of killed brood), a less rigorous measure, is considered biologically significant as the short timeframe for removal should confer disease resistance by removing infected larvae before a pathogen becomes infectious. To assess this, we used liquid nitrogen to freeze an area of sealed brood containing 110 cells. These experimental frames were returned to the respective hives once the nitrogen had volatilized. The amount of dead brood removed by worker bees was measured after 24 hours. For this survey, colonies were categorized as strictly hygienic if 95% or more dead pupae were completely removed from the test area and liberal hygienic if 95% or greater of dead brood was fully or partially removed in 24 hours [33,34].

Hygienic behaviour was calculated as follows:

c= number of cells frozen

u= number of unsealed cells before the test

s= number of sealed cells after the test

p=number of cells partially cleaned out after the test

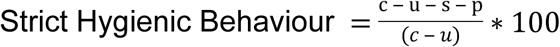

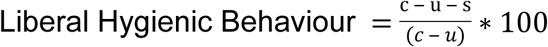

Liberal hygienic behaviour was used to establish a baseline of hygienic behaviour in the colonies sampled. Strict hygienic behaviour was used to assess effectiveness of queen selection in trait transfer, impact of environmental influences, and influence of hygienic behaviour on the control of chalkbrood disease.

## Statistics

### Variables affecting strict hygienic behaviour

Because data were not normally distributed, a binary logistic regression was fitted to the data to understand the relationship between predictors and the likelihood of a colony scoring hygienic or not. The categorical predictor variables: queen selection, breeding program, and nectar flow were considered for the model coded as 0=no and 1=yes. Strict hygienic behaviour was coded as 0= less than 95% removal of dead pupae and 1= greater than 95% removal.

### Nectar flow

Thirty-two colonies were tested twice for hygienic behaviour, once during a nectar flow and once without a nectar flow. The data failed to meet the assumption of sphericity necessary to use a repeated measures test, consequently a nonparametric exact sign test was used to compare the differences in hygienic behaviour with and without a nectar flow. Hygienic behaviour was a continuous variable.

### Chalkbrood infection, nectar flow and hygienic behaviour

A binary logistic regression was fitted to the data to determine if the levels of hygienic behaviour, nectar flow or the interaction of hygienic behaviour and nectar flow were significant predictors of chalkbrood infection. Chalkbrood presence and nectar flow were coded 0=no and 1=yes. Strict test of hygienic behaviour was a continuous variable.

### Severity of Chalkbrood infection and Hygienic behaviour

A Kruskal Wallis test was performed to examine the level of hygienic behaviour in relation to the severity of chalkbrood infection in honey bee colonies. Strict test of hygienic behaviour was a continuous variable and chalkbrood was ordinal with grouping variables: no infection, slight infection, moderate infection, and severe infection. All analyses were carried out in IBM SPSS Statistics for Macintosh, Version 25.0.

## Results

### Colony Summary

Of the 649 colonies tested for hygienic behaviour, 219 (34%) scored above 95% liberal hygienic threshold and 106 (16%) scored above 95% strict hygienic threshold Fig 1. These data can be separated into four categories: no breeding program and no queen selection (35%), breeding program and no queen selection (34%), queen selection without a breeding program (8%), and queen selection only (30%) Table 1.

**Table 1.**
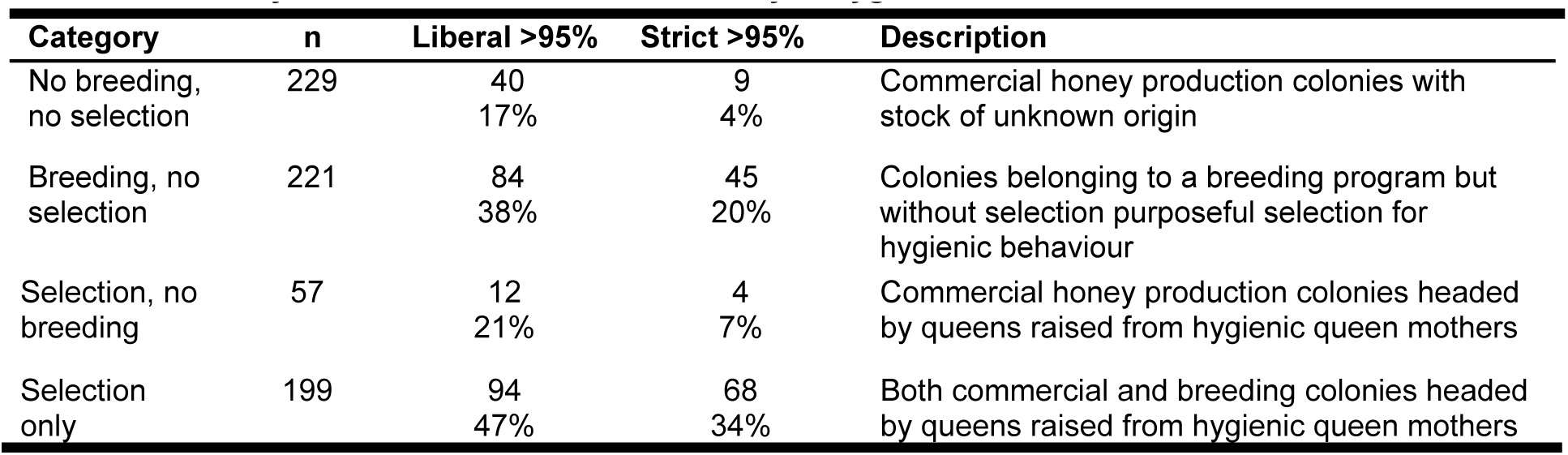
Summary of colonies included in the survey of hygienic behaviour in Australia 2014-2016

| Category | n | Liberal >95% | Strict >95% | Description |
| --- | --- | --- | --- | --- |
| No breeding, no selection | 229 | 40<br>17% | 9<br>4% | Commercial honey production colonies with stock of unknown origin |
| Breeding, no selection | 221 | 84<br>38% | 45<br>20% | Colonies belonging to a breeding program but without selection purposeful selection for hygienic behaviour |
| Selection, no breeding | 57 | 12<br>21% | 4<br>7% | Commercial honey production colonies headed by queens raised from hygienic queen mothers |
| Selection only | 199 | 94<br>47% | 68<br>34% | Both commercial and breeding colonies headed by queens raised from hygienic queen mothers |

**Fig 1.**
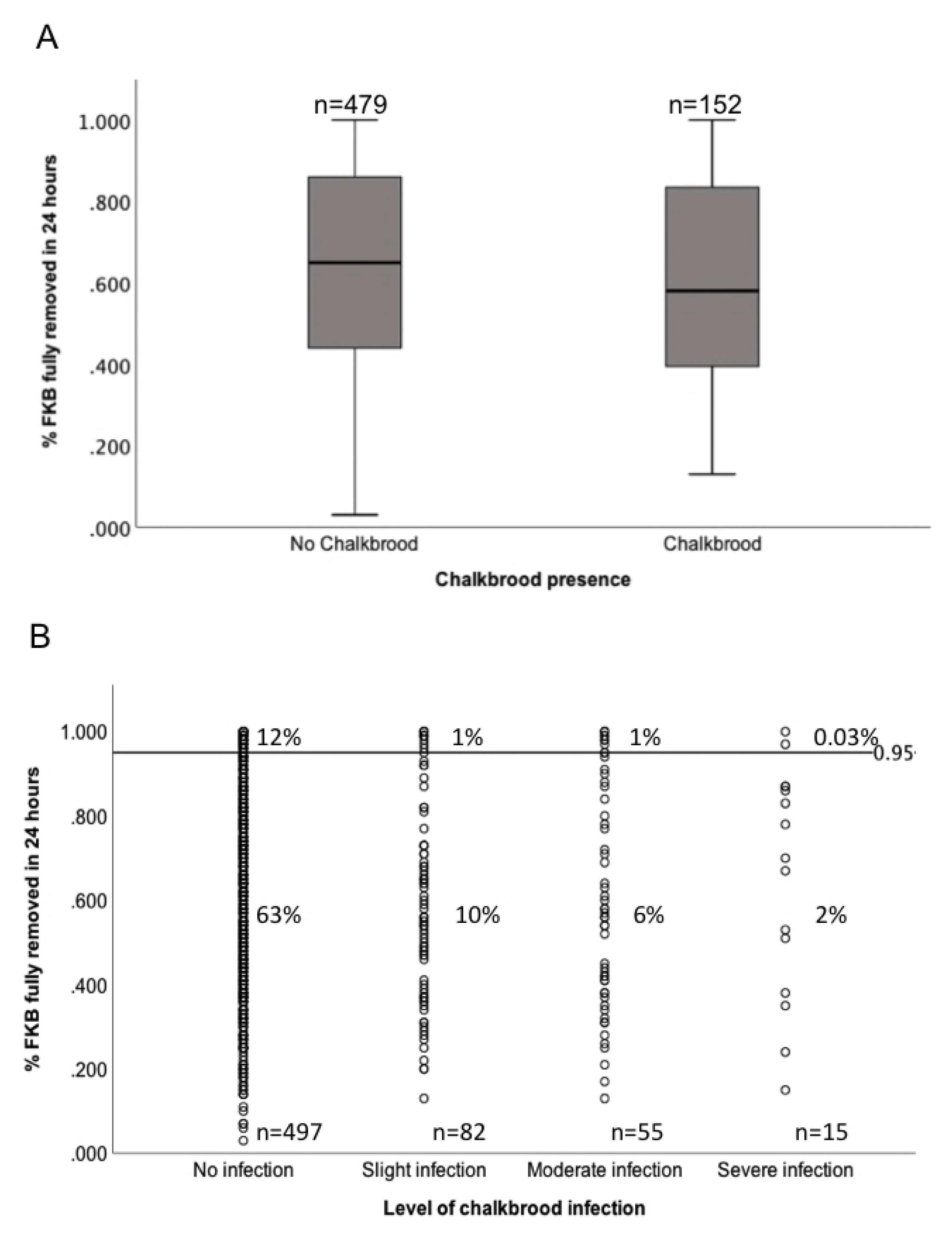
Histograms of Hygienic behaviour in eastern Australian honey bee colonies. Histograms of hygienic behaviour (FKB) removal rates based on the liberal and strict tests from Australian honey bee colonies (n=649) without prior selection for hygienic behaviour from honey production (top) and selective breeding colonies (bottom). Tested from 2014-2016 (A) Strict hygienic behaviour test and (B) Liberal hygienic behaviour test

### Logistic Regression Model

A binary logistic regression analysis was conducted to determine if queen selection, membership of a breeding program, and nectar flow were significant predictors of a colony exhibiting strict hygienic behaviour (<95% removal of FKB in 24 hours) Table 2.

**Table 2.**
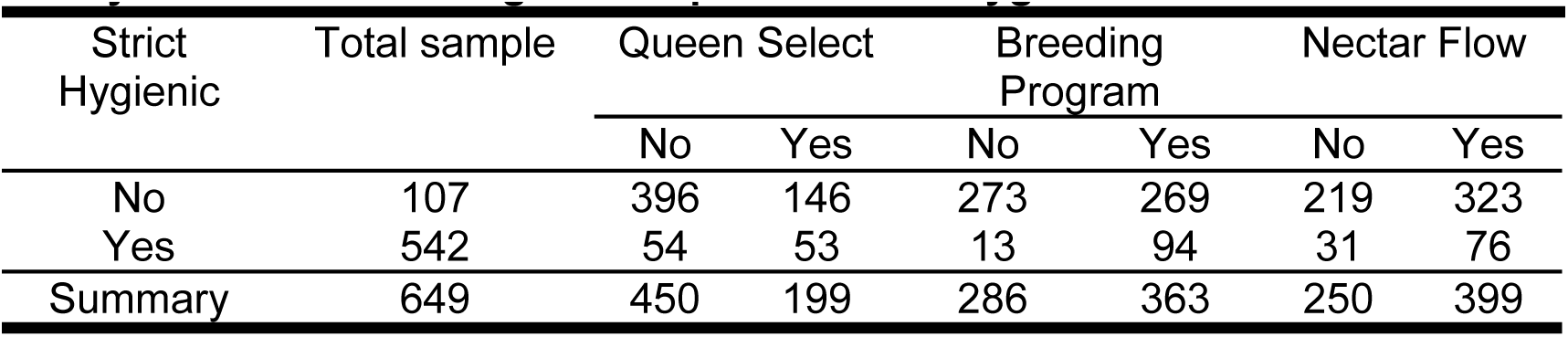
Summary of colony status used in binary logistic regression analysis to determine significant predictors of hygienic behaviour.

A test of the full model against a constant only model was statistically significant, indicating that the predictors reliably distinguished between hygienic and non-hygienic colonies (chi square = 72.550, p < 0.0001 with df = 3) Table 3. Nagelkerke’s R^2^ of 0.179 indicated that this is a weak model explaining 17.9% of the variation. Prediction success overall was 83.5% (100% correctly classified as not strict hygienic and 0% correctly classified as strict hygienic) and was not different from the constant only model indicating that there is a large amount or error in the model. Despite the large error factor, queen selection and breeding program made significant contributions to the predictive capacity of the model (p = 0.002, p < 0.0001, respectively) and can influence a colony’s hygienic status. In fact, the odds ratios indicate that colonies from breeding programs were around 5.8 times more likely to be hygienic and colonies headed by queens raised from hygienic queen mothers were around 2.2 times more likely to be hygienic. Nectar flow was not a significant predictor of strict hygienic behaviour but could none-the-less have a biologically relevant impact on hygienic removal.

**Table 3.** Summary of logistic regression analysis for variables predicting hygienic behaviour (FKB strict test) in Australian honey bee colonies (n=649).

| Observed |  |  | Predicted |  |  |
| --- | --- | --- | --- | --- | --- |
|  |  |  | Not Strict Hygienic | Strict Hygienic | Percentage correct |
|  | Not Strict hygienic |  | 542 | 0 | 100 |
|  | Strict Hygienic |  | 107 | 0 | 0.0 |
| Overall Percentage |  |  |  |  | 83.5 |
| Nagelkerke R<sup>2</sup> |  |  |  |  | 0.719 |

| Variable | B | SE B | Wald | df | P | Odds Ratio |
| --- | --- | --- | --- | --- | --- | --- |
| Queen Selection | 0.818 | 0.237 | 11.903 | 1 | 0.001 | 2.226 |
| Breeding Program | 1.759 | 0.318 | 30.579 | 1 | <0.0001 | 5.807 |
| Nectar Flow | 0.420 | 0.257 | 2.677 | 1 | 0.102 | 1.522 |
| Constant | -3.464 | 0.312 | 109.065 | 1 | <0.0001 | 0.031 |

Dependent variable: strict hygienic behaviour. Predictor variables: queen selection, breeding program, and nectar flow.

### Nectar flow

An exact sign test was used to compare differences in hygienic behaviour when colonies had access to a nectar flow and when they did not. Overall, 32 colonies were tested twice: once during a nectar flow and once without access to nectar. Colonies tested under plentiful nectar conditions had a statistically significant mean increase in strict hygienic behaviour (85.7% +/- 0.183) compared to test scores without nectar access (69.7% +/- 0.256), p= .001, Table 4.

**Table 4.** Descriptive statistics of 32 colonies tested twice for hygienic behaviour: once during a nectar flow and once without a nectar flow.

|  | Mean | Std. Deviation | Minimum | Maximum |
| --- | --- | --- | --- | --- |
| Nectar flow | 0.8574 | 0.18382 | 0.29 | 1.00 |
| No nectar flow | 0.6970 | 0.25633 | 0.03 | 1.00 |

### Chalkbrood disease, nectar flow and strict hygienic behaviour

Of the 649 colonies tested, 76.6% (497) did not show chalkbrood infection, while 23.4% (152) showed clinical symptoms Fig 2A, S1 Fig. A logistic regression analysis was conducted to determine if hygienic behaviour, nectar flow, or the interaction of hygienic behaviour and nectar flow were significant predictors of chalkbrood infection. A test of the full model against a constant only model was not statistically significant, indicating that level of hygienic behaviour, presence of nectar flow or a combination of both were not significant predictors of chalkbrood infection (chi square = 7.012, p =0.072 with df = 3), Table 5, Fig 3, S2 Fig. Data were also analyzed with hygienic behaviour (both liberal and strict tests) as a bivariate predictors with the same results S1 Table.

**Table 5.**
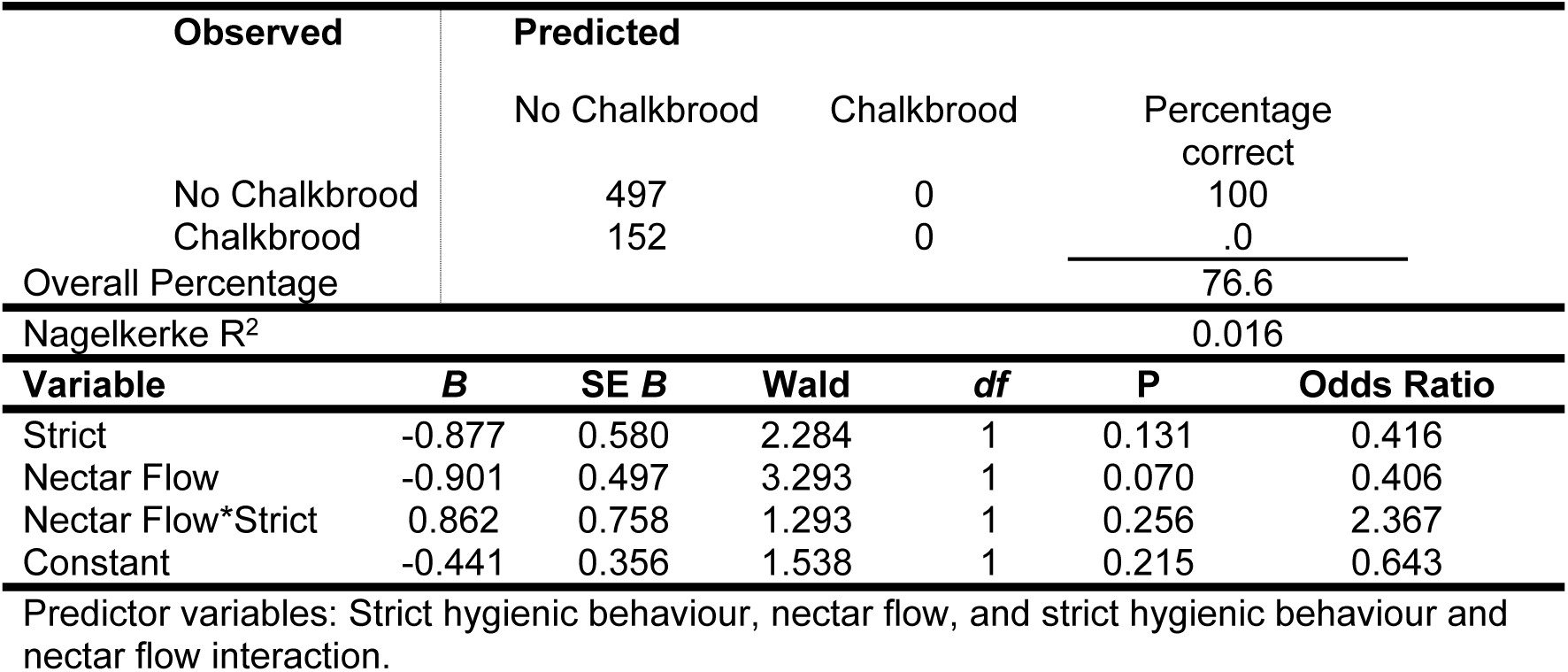
Summary of logistic regression analysis for variables predicting chalkbrood infection. Dependent variable: chalkbrood presence (n=649).

**Fig 2.**
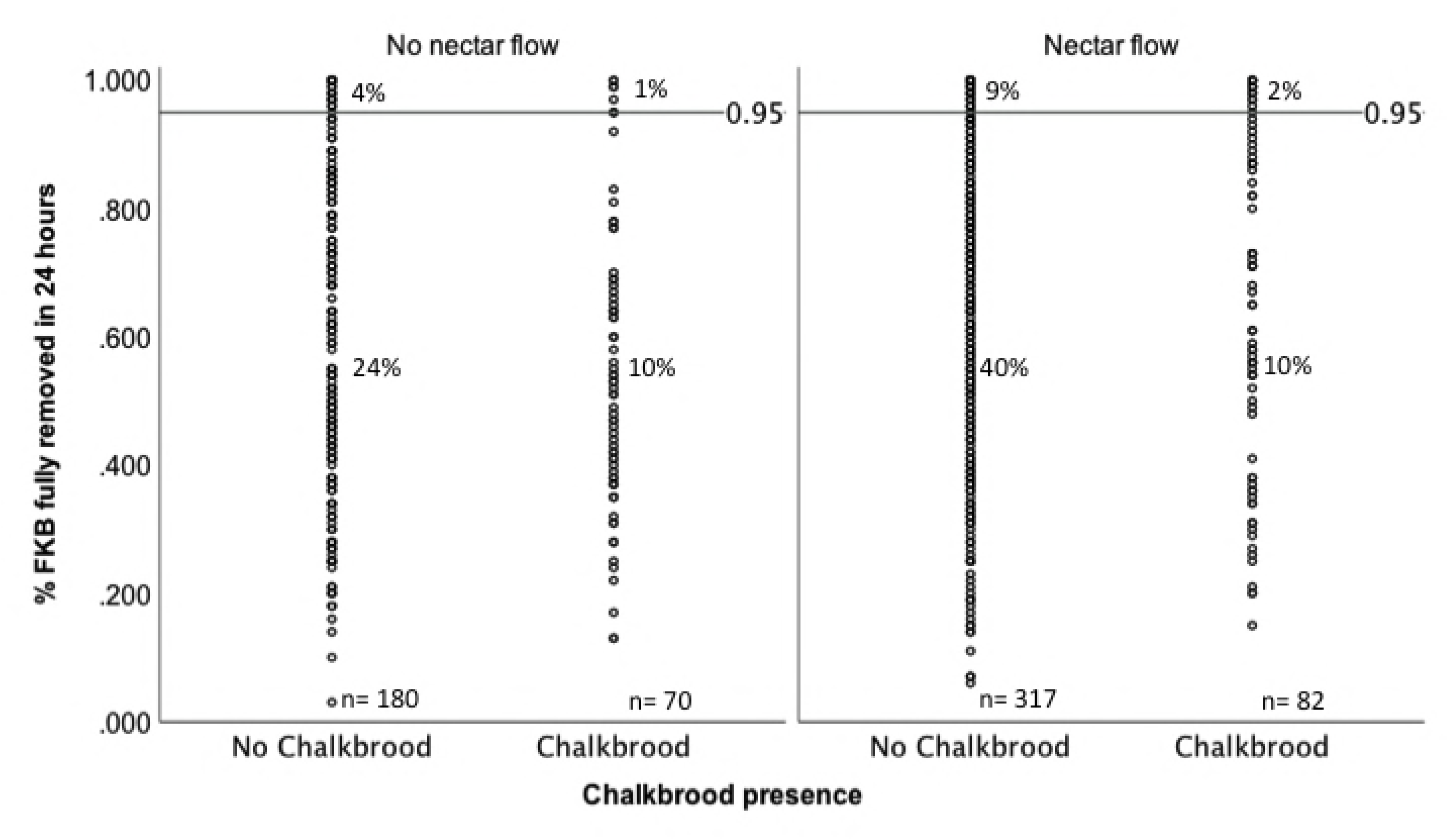
Level of hygienic behaviour based on the FKB strict test not a predictor of chalkbrood infection. (A) Strict hygienic behaviour and chalkbrood infection (B) Severity of chalkbrood infection and strict hygienic behaviour. Percentages are out of 649 colonies

**Fig 3.**
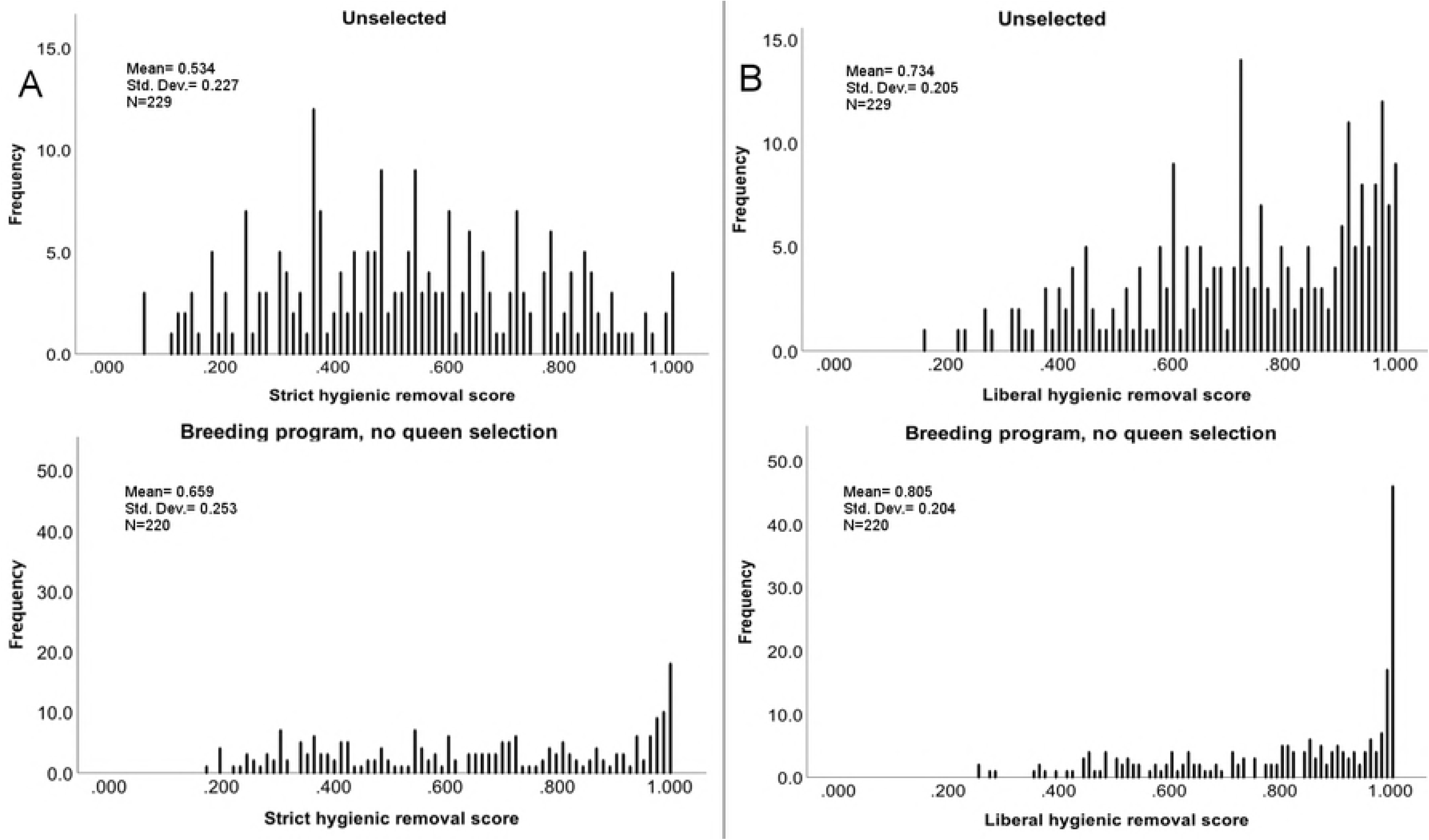
Chalkbrood presence in honey bee colonies not influenced by level of strict hygienic behaviour (FKB test) or nectar flow. Strict hygienic behaviour scores of colonies with and without chalkbrood under variable nectar conditions.

Percentages are out of 649 colonies.

Of the 152 colonies with chalkbrood infection, 53% (82) were slight infections, 37% (55) were moderate infections, and 10% (15) were severe infections Fig 2B. A Kurskal Wallace test determined there was not a statistically significant difference between the level of hygienic behaviour in a colony and absence or level of chalkbrood infection (H(2.461), p=0.482), with a mean rank of 330.79 for no infection, 300.45 for slight infection, 306.51 for moderate infection and 335.17 for severe infection.

## Discussion

The survey aimed to build an understanding of hygienic behaviour in Australian honey bee populations in preparation for selecting and developing hygienic lines of bees given the important contribution of hygienic behaviour to honey bee health. We show that unselected Australian commercial production and breeding populations demonstrate a higher frequency of hygienic colonies (removing <95% frozen brood in 48 hours), 17% and 38% respectively, than similar unselected populations in the Unites States at 10% [26]. These amplified frequencies of hygienic behaviour may be due to directional selection under Australia’s unique floral resource or management practices. Alternatively Australian colonies may be displaying a founder effect from the limited introduction of honey bee genetics from other countries over the years.

Despite the lack of a focused effort to select for and breed hygienic behaviour in Australia, colonies in breeding programs are 5.8 times more likely to be hygienic as per the FKB strict test than those in commercial production operations. These findings indicate that hygienic behaviour may be linked to other economically desirable and heritable traits that enhance a colony’s performance such as general disease resistance or honey production.

This survey found that colonies headed by queens raised from hygienic queen mothers were 2.8 times more likely to be highly hygienic than colonies headed by queens without this type of selection supporting evidence that queen selection alone is sufficient to increase the frequency of this trait in Australian honey bee populations [34,36,37,38]. This may not be the case in populations with a lower frequency of drones originating from hygienic colonies. Spivak and colleagues [34] report that 50% of the drones that hygienic virgin queen mates with must be from hygienic colonies for successful selective breeding.

A degree of trait expression is influenced by nectar availability by driving greater removal of killed brood [34,39,40]. The natural urge of bees to collect nectar and store it as honey may stimulate hygienic behaviour by elevating egg production thereby increasing the need to optimize the use of brood cells. However, Bigio and colleagues [41] report that feeding sugar syrup did not provide the same brood cleaning stimulation. The strict test appears to be more robust than the liberal test providing a good metric for selective breeding purposes regardless of environmental conditions. None-the-less, testing for hygienic behaviour multiple times over the season in periods of both high and low nectar flow will best enable the beekeeper to asses the genetic propensity for removal of dead brood.

The most interesting and surprising component of this survey is that colonies successfully identifying and removing brood killed by liquid nitrogen were not necessarily resistant to chalkbrood infection and the level of hygienic behaviour did not appear to be related to the severity of chalkbrood infection. Many studies demonstrate a relationship between fast removal rates of FKB and reduction of brood disease [9,21,26] but also caution about exclusively relying on this trait for resistance to chalkbrood as fast removal rates of FKB do not always correlate with the removal of chalkbrood infected larvae. Consequently, challenging hygienic colonies with *A. apis* is important for breeding purposes [32] to confirm resistance. To date, most studies that have investigated brood diseases in hygienic colonies have looked at relatively small sample sizes or in experimentally infected colonies. Here we present the first wide-scale survey to verify these cautions.

Hygienic behaviour has been shown to be triggered by phenethyl acetate produced by *A. apis* infected larvae [42]. It is possible that in Australia, chalkbrood infected larvae are not emitting volatiles that initiate hygienic removal. It is unknown what specific biochemical process causes the release of particular volatiles, but if the process is environmentally or nutritionally influenced, eastern Australian conditions and floral resources may not be conducive to the production of important volatiles. Vojvodic and colleagues [43] report variation in virulence between haplotypes of *A. apis.* If the processes involved with volatile production are genetically linked, the lack of important volatiles could result from genetically distinct haplotypes that have responded to directional selection by highly hygienic Australian colonies increasing virulence of specific strains. Yoder and colleagues [44,45] conclude that colony exposure to fungicides can increase chalkbrood disease, but not likely here because our test colonies were mainly surrounded by native flora and not exposed to agrochemicals.

The present work is the first large-scale survey to demonstrate that colonies successfully identifying and removing brood killed by liquid nitrogen are not necessarily resistant to chalkbrood infection in eastern Australia. These results raise the question of whether some specific environmental factor also influences the ability of bees to detect diseased brood and support recommendations that precautions be taken when selecting for hygienic behaviour and chalkbrood resistance [46]. We recommended that the FKB test should be followed up with infection bioassays since the removal of killed brood is not always correlated with the removal of diseases or parasitized brood [32]. However, our recommendation may be problematic because an infection bioassay may not kill all of the larvae since an individual’s innate immunity may be able to overcome the infection [43], making removal rates and social immunity difficult to quantify. Alternatively, the development of an assay to select for social hygienic behaviour specific to eastern Australia’s honey bee population and pest and disease matrix may be beneficial.

Pest and disease resistance is the impetus of breeding for hygienic behavior. Bees bred for Varroa Sensitive Hygiene in other countries were shown to be highly hygienic using the FKB assay even though the population was selected by other means [47,48]. We also show that bees selected for performance (eg honey production, pollination efficacy, temperament, etc) can also be quite hygienic. It is possible, therefore, for many different selection assays and methods to increase the nest cleaning ability of honey bees to remove parasitized brood and thus arrive at the desired outcome: healthier, more disease resistant bees.

The level of hygienic behaviour found in Australia’s managed honey bee colonies is surprising and encouraging, laying a solid framework for selective breeding programs targeting disease resistance and ultimately preparing for living with *Varroa*. However, to maximize the benefits of this important trait for the honey bee industry, a deeper understanding of trait selection and the evolutionary arms race between *A. mellifera* and pathogen analogues such as *A. apis* is vital.

## Acknowledgements

We thank the Australian Honey Bee Industry Council (AHBIC) for supporting the concept of this work and participating beekeepers for their time, expertise, and providing access to their bees. We would specifically like to thank Thomas Harding for his technical beekeeping support. Thank you to Daniel Martin for supporting this work. Thank you to Dr. Graeme Byrne of La Trobe University for statistics assistance and gratitude to our reviewers.

## Supporting information

**S1 Fig. Hygienic behaviour as determined by the liberal FKB test not a predictor of chalkbrood infection** (A) hygienic behaviour and chalkbrood infection (B) Severity of chalkbrood infection and liberal hygienic behaviour. Percentages are out of 649 colonies.

**S2 Fig. Chalkbrood presence in honey bee colonies not influenced by liberal FKB test of hygienic behaviour or nectar flow** hygienic behaviour scores from colonies with and without chalkbrood under variable nectar conditions.

**S3** Data associated with this survey

**S1 Table. Summary of logistic regression analysis for variables predicting chalkbrood infection**

